# Polymeric Micellar Nanoparticles Enable Image-guided Drug Delivery in Solid Tumors

**DOI:** 10.1101/2024.06.07.598019

**Authors:** Jashim Uddin, Justin Han-Je Lo, Mukesh K. Gupta, Thomas A. Werfel, Abu Asaduzzaman, Connor G. Oltman, Eva F. Gbur, Mohammed T. Mohyuddin, Farhana Nazmin, Saidur Rahman, Ahan Jashim, Brenda C. Crews, Philip J. Kingsley, Lawrence J. Marnett, Craig L. Duvall, Rebecca S. Cook

## Abstract

We report the development of a nanotechnology to co-deliver chemocoxib A with a reactive oxygen species (ROS)-activatable and COX-2 targeted pro-fluorescent probe, fluorocoxib Q (FQ) enabling real time visualization of COX-2 and CA drug delivery into solid cancers, using a di-block PPS_135_-*b*-POEGA_17_ copolymer, selected for its intrinsic responsiveness to elevated reactive oxygen species (ROS), a key trait of the tumor microenvironment. FQ and CA were synthesized independently, then co-encapsulated within micellar PPS_135_-*b*-POEGA_17_ co-polymeric nanoparticles (FQ-CA-NPs), and were assessed for cargo concentration, hydrodynamic diameter, zeta potential, and ROS-dependent cargo release. The uptake of FQ-CA-NPs in mouse mammary cancer cells and cargo release was assessed by fluorescence microscopy. Intravenous delivery of FQ-CA-NPs to mice harboring orthotopic mammary tumors, followed by vital optimal imaging, was used to assess delivery to tumors *in vivo*. The CA-FQ-NPs exhibited a hydrodynamic diameter of 109.2 ± 4.1 nm and a zeta potential (ζ) of –1.59 ± 0.3 mV. Fluorescence microscopy showed ROS-dependent cargo release by FQ-CA-NPs in 4T1 cells, decreasing growth of 4T1 breast cancer cells, but not affecting growth of primary human mammary epithelial cells (HMECs). NP-derived fluorescence was detected in mammary tumors, but not in healthy organs. Tumor LC-MS/MS analysis identified both CA (2.38 nmol/g tumor tissue) and FQ (0.115 nmol/g tumor tissue), confirming the FQ-mediated image guidance of CA delivery in solid tumors. Thus, co-encapsulation of FQ and CA into micellar nanoparticles (FQ-CA-NPs) enabled ROS-sensitive drug release and COX-2-targeted visualization of solid tumors.

## INTRODUCTION

Nanomedicine formulations of systemically dosed chemotherapeutic drugs aim to improve biodistribution and target site accumulation of cytotoxic drugs.^1^ Various passively and actively targeted nanomedicines have been evaluated, e.g., liposomes,^2^ polymer micelles,^3, 4^ and antibodies.^5^ Abundant preclinical data show that 5-200 nm sized nanoparticles can improve the therapeutic index of low molecular weight drugs.^1, 2, 6-8^ Nanomedicine formulations also have imaging applications.^4, 9, 10^ Personalized medicine will benefit from a new paradigm that enables co-delivery of imaging and therapeutic agents into cancers for imaging during treatment, complementing current tumor imaging studies that occur only before and after treatment.

Studies have shown show that cyclooxygenase-2 (COX-2) contributes to several pathologies, including cancer.^11^ Importantly, COX-2 and reactive oxygen species (ROS) co-exist in pathogenesis.^12^ COX-2 is an attractive molecular target for delivering imaging/therapeutic agents to tumors because it is expressed in only a few normal tissues and is greatly up-regulated in inflamed tissues and many premalignant and malignant tumors. Radiolabeled COX-2 inhibitors have been developed for nuclear imaging. We reported the synthesis and *in vivo* validation of radiologic imaging agents for both SPECT and PET imaging. The structures of these agents are based on the indomethacin and celecoxib scaffolds. We validated the specificity and sensitivity of our probes in COX-2-targeted imaging of inflammation and cancer both *in vitro* and *in vivo, and* in *ex vivo*. In addition, fluorescent COX-2 inhibitors are attractive candidates as targeted optical imaging agents. Such compounds have the advantage that each molecule bears the fluorescent labeling, and the compounds are non-radioactive and stable. Thus, they can be used for cellular imaging, animal imaging, and clinical imaging of tissues where topical or endoluminal illumination is possible (e.g., esophagus, colon, and upper airway via endoscopy, colonoscopy, and bronchoscopy, respectively). Prior work from our laboratory demonstrated that fluorescent COX-2 inhibitors are useful as chemical probes for protein binding and *in vivo* imaging. We designed and synthesized COX-2-targeted imaging agents and later we successfully discovered a novel redox-activatable COX-2-targeted optical imaging agent, called fluorocoxib Q. The fluorocoxib Q is a nitroxide derivative of fluorocoxib A. Fluorocoxib Q enables targeted visualization of COX-2 and ROS in pathological tissues. Fluorocoxib Q exhibits extremely low fluorescence emission due to quenching of the excited electronic state of the carboxy-X-rhodamine by the nitroxide radical within the molecule. Upon radical trapping in cancer cells by reacting oxygen species (ROS), the COX-2-targeted fluorocoxib Q probe becomes fluorescently activated, making it effective for tumors imaging.

Overexpression of COX-2 and its prostaglandin products play a vital role in tumorigenesis. Although enormous research has been conducted, the relative importance of each of these effects is still unknown. Several clinical trials have been completed and many are ongoing to evaluate specific COX-2 inhibitors for cancer chemoprevention, only celecoxib has been approved by FDA for the treatment of familial adenomatous polyposis (FAP). Our prior studies demonstrate COX-2-targeted delivery of imaging agents to tumors,^9, 11, 13-19^ suggesting the possibility that this approach can be used to selectively deliver chemotherapeutic agents as well. An attempt to test this hypothesis we used a conjugate chemistry approach, where we discovered a cytotoxic COX-2 inhibitor, called chemocoxib A ^20^, a conjugate of indomethacin with podophyllotoxin, revealed highly potent and selective COX-2 inhibition in purified enzyme and in cell-based assays.

X-ray co-crystallographic studies demonstrated the structural basis of COX-2 binding by chemocoxib A and kinetic analysis demonstrated that chemocoxib A is a slow, tight-binding inhibitor of COX-2.^21^ The conjugate exhibited cytotoxicity in cell culture like that of podophyllotoxin. It was accumulated selectively in COX-2 positive tumors in mice and induced tumor growth inhibition, but didn’t have any adverse effect on tumors that did not express COX-2 enzyme.^21^ It should be noted that podophyllotoxin itself does not inhibit COX-2, but when tethered to indomethacin it does, and accumulates in COX-2expressing tumor cells. How it slows growth of cancer cells *in vitro* and *in vivo* is not fully understood, but it does not affect the viability of primary human mammary epithelial cells (HMECs) in culture. Also, it does not cause systemic toxicity in animals.^21^ Thus, chemocoxib A provides a proof-of-concept for *in vivo* targeting of chemotherapeutic agents to COX-2 and represents the 1^st^ cytotoxic COX-2 inhibitor validated for targeted tumor growth inhibition *in vivo*. We hypothesize that fluorocoxib Q^12^ and chemocoxib A^21^ can be co-encapsulated into ROS-responsive polymeric micellar nanoparticles that can codeliver both agents into tumors allowing image-guided drug delivery into the solid tumors with molecular specificity in real time.

Herein, we report the discovery of FQ and CA co-loaded PPS_135_-*b*-POEGA_17_ polymeric micellar nanoparticles (FQ-CA-NPs) and their delivery into COX-2 expressing breast cancer cells and orthotopic mouse mammary tumors. By combining disease diagnosis and therapy in one medical specimen, this study resulted in a new nanoplatform, where targeted tumor codelivery of imaging and therapeutic agents has been achieved.

## MATERIALS AND METHODS

### Chemical Synthesis

Fluorocoxib Q (FQ), chemocoxib A, and the di-block PPS_135_-*b*-POEGA_17_ copolymer were synthesized using previously published procedures.^9, 12, 21, 22^

### Co-encapsulation of FQ and CA in PS_135_-*b*-POEGA_17_ Polymeric Micellar Nanoparticles

FQ and CA were co-encapsulated into PPS_135_-*b*-POEGA_17_ copolymeric micelles using a bulk solvent evaporation method.^9^ At first, FQ, CA, and PPS_135_-*b*-POEGA_17_ were each dissolved in chloroform separately. The FQ (10 mL, 20 mg/mL) and CA (10 mL, 20 mg/mL) solutions were added to the PPS_135_-*b*-POEGA_17_ copolymer (20 mL, 200 mg/mL) solution. The resultant solution was added dropwise to a Dulbecco’s Phosphate Buffered Saline (1 mL, DPBS pH 7.0– 7.4, Ca^2+^ and Mg^2+^-free) followed by 16 h gentle stirring at 25°C in the dark. A sterile centrifugation of the resultant aqueous solution afforded FQ and CA co-loaded nanoparticles (FQ-CA-NPs) ready for *in vitro* and *in vivo* evaluations.

### Concentration of FQ and CA in Micellar Nanoparticles

Concentration of CA and FQ within micellar nanoparticles (FQ-CA-NPs, 0.5 mL in DPBS) was measured by first disassembling NPs in dimethylformamide (DMF, 0.5 mL, stirring 16 h, at 25°C) Samples were dried, reconstituted and analyzed by reversed phase HPLC-UV using Phenomenex C18 columns held at 40°C. Each compound was quantified against a standard curve obtained for each after dissolving them in 50% DMF in DPBS. The statistical comparisons of the experimental results were performed by Student’s t-test at a significance level of 0.01 and 0.001, which afforded FQ and CA concentrations to be 0.132 mg/mL and 0.147 mg/mL in a typical FQ-CA-NPs injectable formulation.

### Properties of FQ-CA-NPs

A Malvern Zetasizer Nano-ZS Instrument equipped with a 4 mW Helium Neon Laser operating at 632.8 nm was used to measure the hydrodynamic diameter (D_h_) and zeta potential (ζ) of the micellar FQ-CA-NPs.

### H_2_O_2_-Dependent Cargo Release from FQ-CA-NPs

FQ-CA-NPs were incubated in H_2_O_2_ (ROS) at progressively increasing concentrations (0 to 1000 mM) in 96-well plates. Fluorescence intensity was monitored using a Tecan Infinite 500 plate reader (excitation 540 nm, emission 610 nm), normalizing each sample to its fluorescence value prior to addition of H_2_O_2_.

### Fluorescence Microscopy of 4T1 Cells

Mouse 4T1 mammary carcinoma tumor cells were obtained from the Cell Culture and Tissue Engineering Core of the Vanderbilt Institute for Integrative Biosystems Research and Education. The 4T1 cells (1.6 × 10^5^ cells) were plated on MatTek glass-bottom culture dishes. The 4T1 Cells in Hank’s balanced salt solution (HBSS)/Tyrode’s were grown to a 60% confluency and incubated with 200 nM of fluorocoxib Q for 3 h at 37 °C. Adherent cells were treated with FQ-CA-NPs (1 μM). Cells were rinsed 3 times with Hanks Balanced Salt Solution (HBSS)/Tyrode’s buffer and imaged at a gain of 20 on a Leica DM IL LED FIM fluorescence microscope.

### Cell Viability Assay

Primary human mammary epithelial cells (HMECs) were obtained as a gift from Dr. Jennifer Pietenpol, Vanderbilt University Medical Center, and mouse 4T1 mammary carcinoma cells were obtained from the Cell Culture and Tissue Engineering Core of the Vanderbilt Institute for Integrative Biosystems Research and Education. HMECs were grown in HMEC medium containing Mammary Epithelial Growth Supplement (Life Technologies) or mouse 4T1 mammary carcinoma cells were grown in growth media (Dulbecco’s Modified Eagle Medium:Nutrient Mixture F12 plus 10% fetal bovine serum and antibiotic/antimycotic). Cells (8,000 to 10,000/well) were plated in 96-well plates (Sarstedt). After 24 hours, cells were treated with FQ-CA-NPs (ca. 200 nmol of CA) or vehicle in fresh growth media. Viability was assessed 48 hours later using WST-1 Cell Proliferation Reagent (Roche, 11644807001) as described previously.^23^

### Vital Fluorescence Imaging and Organ Biodistribution in a Mouse Model of Orthotopic Breast Cancer

All experiments with mice were approved by the Institutional Animal Care and Use Committees (IACUC) at the Vanderbilt University School of Medicine. 4T1 cells (1 × 10^6^ cells) were inoculated into the left inguinal mammary fatpad.^24^ Tumors were measured twice weekly until reaching 700-900 mm^3^. CA-FQ-NPs (1 mg/kg of each CA and FQ) was delivered by subcutaneous injection. At 49 h post-injection, we lightly anesthetized dosed animals with 2% isoflurane and imaged them using a Xenogen IVIS 200 Optical Imaging Instrument with a DsRed filter spanning 570–615 nm with a 20 cm field of view at 20 microns resolution at a depth of 1.5 cm and an exposure of 1 sec. Using an ImageJ software, we analyzed the static images obtained from the optical imaging of tumor bearing animals (n = 6 animals/group), where regions of interest (ROIs) were created for measurement. After *in vivo* imaging, mice were humanely euthanized and tumor, brain, liver, lung, and kidney were collected. The dissected organs were imaged on a Xenogen IVIS 200 Optical Imaging Instrument. Fluorescence intensity (photons/sec) in each static image was measured using an ImageJ software. We determined the signal-to-noise ratios by dividing the mean fluorescence intensity of tumorous breast (1.25e+009, n = 6 tumors) with the mean fluorescence intensity of celecoxib (10 mg/kg dose)-pretreated breast (1.86e+007, n = 6 tumors) within the respective ROIs measured in photons/sec using an ImageJ software.

### Statistical Methods

Student’s t-test was used for statistical analyses of fluorescence signal intensities in static images of breast tumors or major organs including lung, liver, kidney, or brain, where statistical significance at *P* ≤ 0.05 was considered. Within the size of the samples (*n*), signal intensities were used as the arithmetic mean and standard error.

## RESULTS AND DISCUSSION

We synthesized fluorocoxib Q (FQ),^12^ a nitroxide analog of fluorocoxib A.^11^ FQ is pro-fluorescent, which is due to quenching of its carboxy-X-rhodamine fluorescence by the nitroxide radical within the molecule and which is a redox-activatable COX-2-taregeted imaging agent exhibiting extremely low fluorescence emission until activated by ROS present in pathologic tissues.^12^ Next, we synthesized chemocoxib A ^20^, a cytotoxic COX-2 inhibitor exhibiting potent antitumor activity.^21^ CA is an indomethacin–podophyllotoxin conjugate demonstrated tumor growth inhibition without any systemic toxicity.^21^

In addition, we synthesized PPS_135_-*b*-POEGA_17_, an amphiphilic di-block co-polymer that self-assembles into micellar nanoparticles <200 nm in hydrodynamic diameter.^9^ Then, we co-loaded FQ and CA into ROS-responsive micellar nanoparticles (FQ-CA-NPs) of a di-block copolymer (PPS_135_-*b*-POEGA_17_) (Figure 1) using a bulk solvent evaporation method.^9^ In this method, FQ, CA and PPS_135_-*b*-POEGA_17_ were dissolved in chloroform singly. Then, FQ and CA solutions were mixed, and the mixture was added to the PPS_135_-*b*-POEGA_17_ copolymer solution, then the resultant solution was added dropwise to 1 mL of Dulbecco’s phosphate buffered saline (DPBS pH 7.0 to 7.3, without calcium and magnesium) at 25°C with gentle stirring. The binary system was stirred for 16 h at 25°C in the dark, during which time the chloroform was evaporated. Sterile centrifugation of the formulation afforded FQ and CA co-loaded micellar nanoparticles (FQ-CA-NPs) ready to be used for evaluation and visualization of cells and CA delivery in orthotopic mammary tumors implanted in nude mice.

**Figure 1.**
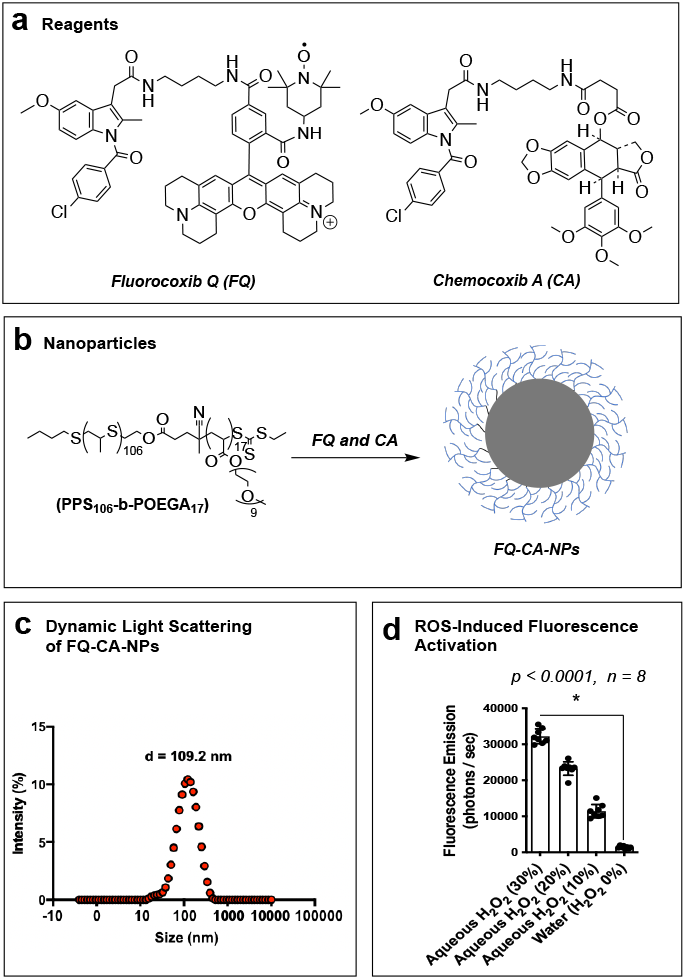
(a) Chemical structure of fluorocoxib Q (FQ) and chemocoxib A^20^. (b) Co-encapsulation of FQ and CA using a bulk evaporation method for the formation of micellar nanoparticles (FQ-CA-NPs) of PPS_135_-*b*-POEGA_17_ in a 50:50 mixture of CHCl_3_/DPBS (v/v) with a gentle stirring at 25°C for 16 h. (c) Dynamic light scattering of FQ-CA-NPs. (d) Light emission from FQ-CA-NPs treated with aqueous hydrogen peroxide (H_2_O_2_) solutions.

We determined the hydrodynamic diameter of FQ-CA-NPs to be 109.2 ± 4.1 nm and a zeta potential (ζ) of –1.59 ± 0.3 mV. We treated FQ-CA-NPs with H_2_O_2_ solutions at a range concentration that showed higher fluorescence with 30% as compared to 20% or lower H_2_O_2_ concentration in water, suggesting its sensitivity to ROS allowing drug release and fluorescence activation of FQ. We evaluated the co-loaded nanoparticles in optical imaging of cells, where FQ-CA-NPs showed higher fluorescence in vehicle-pretreated 4T1 cells as compared to celecoxib-pretreated 4T1 cells (Figure 2a) with a significant difference (Figure 2b). In a cell viability assay, the FQ-CA-NPs was toxic to 4T1 breast carcinoma cells with an EC_50_ of 1.66 μM, but nontoxic to primary human mammary epithelial cells (HMECs). (Figure 2c).

**Figure 2.**
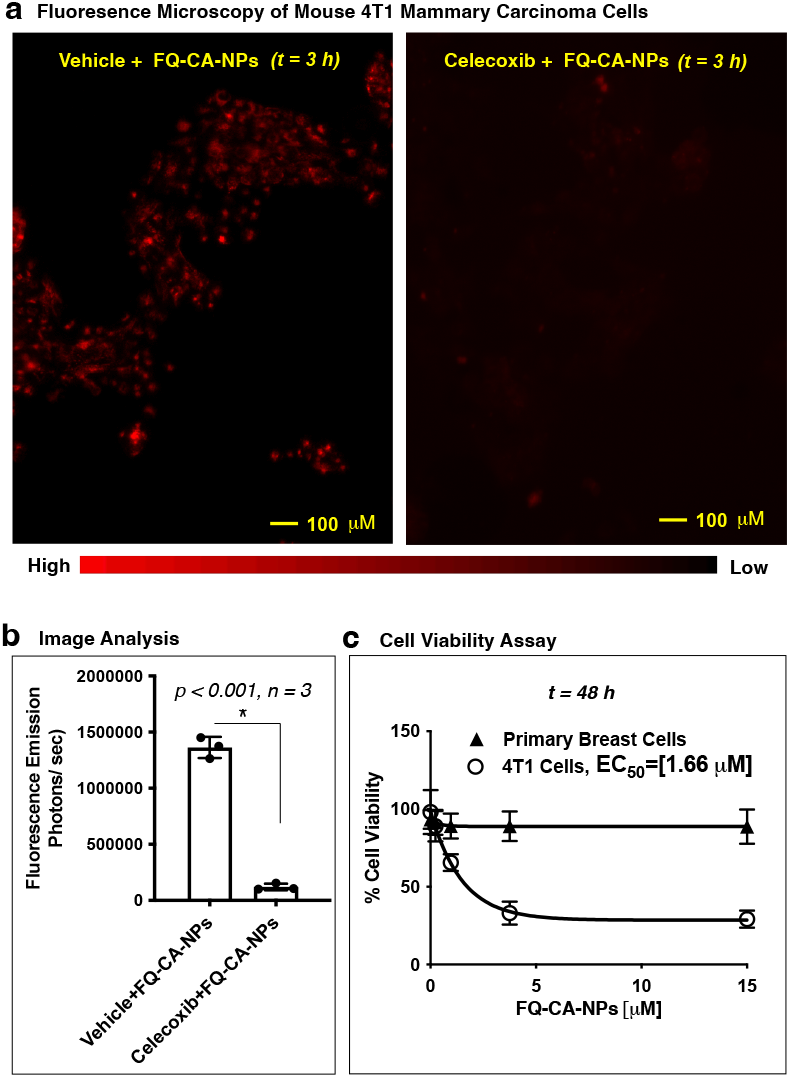
(a) Fluorescence microscopy of vehicle or celecoxib pre-treated 4T1 cells incubated FQ-CA-NPs for 3 h. (b) Image analysis of vehicle or celecoxib pretreated 4T1 cells incubated with FQ-CA-NPs using an ImageJ software. (c) Viability of primary human mammary epithelial cells (HMECs) or 4T1 mouse mammary carcinoma cells treated with FQ-CA-NPs for 48 h.

We determined the *in vivo* pharmacokinetics of FQ and CA in C57BL/6 mice. In this assay, FQ resulted in a plasma half-life of 26.4 h and CA showed 9.3 h (Figure 3a). We dosed FQ-CA-NPs by subcutaneous injection to mice harboring orthotopic mouse mammary tumors (4T1) expressing COX-2 enzyme.^25^ Optical imaging (Xenogen IVIS200) at 49 h post-injection detected a high fluorescence in the tumor to suggest the uptake of FQ-CA-NPs and ROS-mediated release of FQ and CA in the tumor followed by fluorescence activation of FQ (non-fluorescent) to FQ-H (fluorescent) (Figure 3b-c). Fluorescence signal was lowered significantly in the mammary tumor of the animal pre-dosed with 10 mg/kg (i.p.) of tempol or celecoxib followed by FQ-CA-NPs injection. The tempol pre-treatment traps tumor ROS to prevent cargo release from the loaded-NPs and celecoxib pre-treatment pre-clued binding of both FQ or CA with COX-2 by its active site blockade. After imaging, the animals were euthanized and the tumor, brain, liver, lung, and kidney were collected and imaged under Xenogen IVIS200 camera to measure the fluorescence intensity in these tissues. The measurement showed significantly higher tumor signal in FQ-CA-NPs animals compared to tempol or celecoxib pre-treated animal tumors (Figure 3d-e).

**Figure 3.**
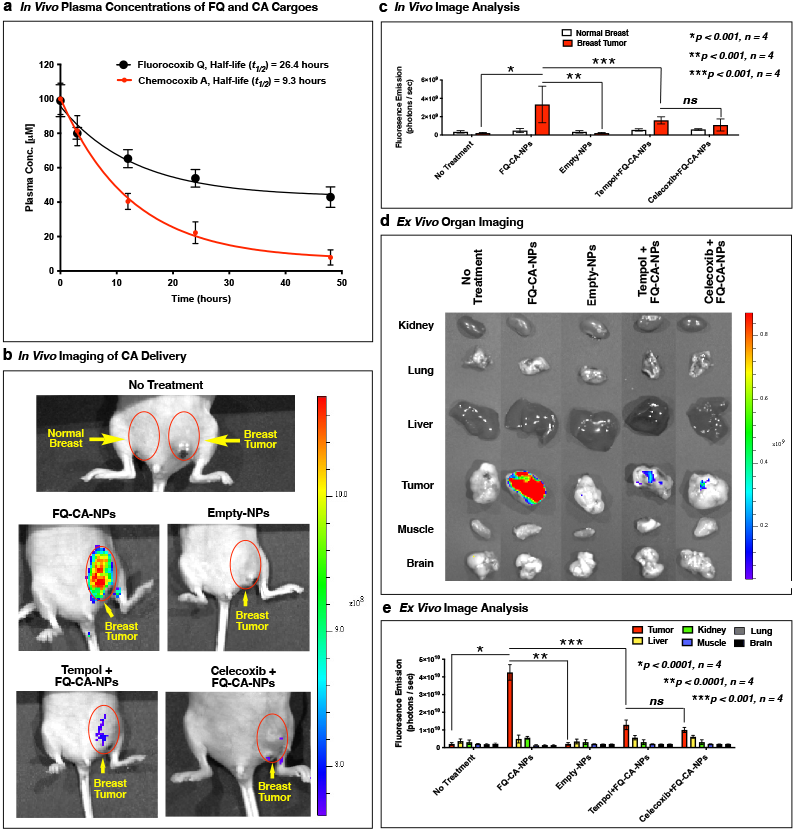
(a) *In vivo* pharmacokinetics of FQ and CA in C57BL/6 mice. (b) *In vivo* optical imaging of NU/J mice bearing orthotopic 4T1 mammary tumors at 49 h post-dosing subcutaneous administration of FQ-CA-NPs, Empty-NPs, Tempo followed by FQ-CA-NPs, and celecoxib followed by FQ-CA-NPs. (c) Measurement of light emission in the normal breast and breast tumors by an ImageJ software. (d) *Ex vivo* optical imaging of kidney, lung, liver, tumor, muscle and brain of NU/J mice bearing orthotopic 4T1 mammary tumors at 49 h post-dosing intravenous administration of FQ-CA-NPs, Empty-NPs, Tempo followed by FQ-CA-NPs, and celecoxib followed by FQ-CA-NPs. (e) Measurement of light emission in the excised kidney, lung, liver, tumor, muscle and brain by an ImageJ software.

To confirm the uptake of FQ-CA-NPs, the tumor tissues were analyzed by LC-MS/MS, which quantified the amount of FQ (0.115 nmol/g tissue) and CA (2.38 nmol/g tissue) delivered, released and retained in the tumor. The fluorescence imaging and the bioanalytical data confirmed the delivery of FQ-CA-NPs followed by FQ-mediated visualization of CA-delivery in the mammary tumor.

These studies showed the feasibility of *in vivo* targeting of COX-2 in tumors using co-loaded micellar nanoparticles to display delivery of loaded compounds into the breast tumors in live animals. In presence of ROS in the breast tumors, the micellar nanoparticles (FQ-CA-NPs) were destabilized such that release of both cytotoxic agent CA and activatable probe FQ in tumor cells can take place to bind with intracellular COX-2 for enhanced tumor retention, where the activatable COX-2 probe FQ became fluorescently activated by tumor ROS allowing visualization of tumor delivery of cytotoxic agent ^20^ in real-time and fluorescence activation of FQ was possible by interaction with ROS, co-localized with COX-2 at the tumor site.^12^ In addition, the experimental results suggests that the tumor uptake of FQ-CA-NPs followed by ROS-induced nanoparticle disassembly and release of encapsulated compounds for activation FQ into the fluorescent species for COX-2 binding to produce a detectable buildup of fluorescence signal required longer time (49 h) as compared to the free FQ compound (24 h).^12^

## CONCLUSIONS

We report the discovery and characterization of FQ-CA-NPs, a nanomedicine formulation based on polymeric micellar nanoparticles of FQ and CA. The FQ-CA-NPs showed sensitivity to ROS. Treatment of FQ-CA-NPs with vehicle-pretreated 4T1 cells showed higher fluorescence compared to celecoxib-pretreated 4T1 cells. FQ-CA-NPs showed toxicity to 4T1 breast carcinoma cells but is non-toxic to primary mammary cells. Following systemic administration, FQ-CA-NPs enabled co-delivery of FQ and CA in orthotopic mammary tumors. Fluorescence activation of FQ in the tumor microenvironment allowed its fluorescence activation and COX-2 binding for tumor accumulation and retention, and detection of CA delivery into solid mammary tumors. The targeted co-delivery of FQ and CA was confirmed by LC-MS/MS analysis of tumor tissues.

## Supporting information

Supporting Information BioRxiv1

## CORRESPONDING AUTHORS

*Md. Jashim Uddin, Ph.D., phone: 615-484-8674, and Rebecca S. Cook, Ph.D.

## AUTHOR CONTRIBUTIONS

M.J.U. conceived the idea, synthesized fluorocoxib Q and chemocoxib A, designed the experiments, performed the polymeric micellar nano-formulation, conducted animal imaging experiments, acquired and interpreted imaging and analytical data, wrote the manuscript, and supervised the overall conduct of the research program; C.G.O., J.H.L., and M.M. assisted M.J.U. in the synthesis and chromatographic purification of fluorocoxib Q and chemocoxib A; M.K.G. synthesized the polymer; T.A.W. assisted M.J.U. in nano-formulation and characterization of co-loaded micellar nanoparticles; F.N. and M.S.R. performed the image analysis; B.C.C. performed the cell imaging and viability assays; P.J.K. performed tissue analysis using LC-MS/MS; C.L.D. and L.J.M provided laboratory space, and R.S.C. developed and provided the mouse model of orthotopic mammary tumor.

## Personnel and Facilities

The authors are grateful to Dr. Dmitry S. Koktysh of the Vanderbilt Institute of Nanoscale Science and Engineering Core Facility for assistance with transmission electron microscopy in measuring hydrodynamic diameter and transmission electron microscopy of the co-loaded nanoparticles; the Vanderbilt University Institute of Imaging Sciences for the use of the Xenogen IVIS 200 imaging system, the Vanderbilt Small Molecule NMR Facility for the use of the Bruker AV-I at 600 MHz instrument, and the Vanderbilt Mass Spectroscopy Research Center for use of the Quantum Triple Quadrupole instrument.

### Funding Sources

This program has been financially supported by research grants from the Phi Beta Psi Sorority Trust AWD00000652 and AWD00001248 to M.J.U.; the Vanderbilt Institute for Clinical and Translational Research VR52653 to M.J.U.; and the National Institutes of Health R01 CA89450 to L.J.M., and R01 CA260958-01A1 to M.J.U., C.L.D., & R.S.C.

## REFERENCES

(1) Sofias, A. M.; Dunne, M.; Storm, G.; Allen, C. The battle of “nano” paclitaxel. Adv Drug Deliv Rev 2017, 122, 20–30. DOI: 10.1016/j.addr.2017.02.003.

(2) Allen, T. M.; Cullis, P. R. Liposomal drug delivery systems: from concept to clinical applications. Adv Drug Deliv Rev 2013, 65 (1), 36–48. DOI: 10.1016/j.addr.2012.09.037.

(3) Ghezzi, M.; Pescina, S.; Delledonne, A.; Ferraboschi, I.; Sissa, C.; Terenziani, F.; Remiro, P. F. R.; Santi, P.; Nicoli, S. Improvement of Imiquimod Solubilization and Skin Retention via TPGS Micelles: Exploiting the Co-Solubilizing Effect of Oleic Acid. Pharmaceutics 2021, 13 (9). DOI: 10.3390/pharmaceutics13091476.

(4) Ghezzi, M.; Pescina, S.; Padula, C.; Santi, P.; Del Favero, E.; Cantu, L.; Nicoli, S. Polymeric micelles in drug delivery: An insight of the techniques for their characterization and assessment in biorelevant conditions. J Control Release 2021, 332, 312–336. DOI: 10.1016/j.jconrel.2021.02.031.

(5) Alley, S. C.; Okeley, N. M.; Senter, P. D. Antibody-drug conjugates: targeted drug delivery for cancer. Curr Opin Chem Biol 2010, 14 (4), 529–537. DOI: 10.1016/j.cbpa.2010.06.170.

(6) Kotta, S.; Aldawsari, H. M.; Badr-Eldin, S. M.; Nair, A. B.; Yt, K. Progress in Polymeric Micelles for Drug Delivery Applications. Pharmaceutics 2022, 14 (8). DOI: 10.3390/pharmaceutics14081636.

(7) Sharma, S.; Pukale, S. S.; Sahel, D. K.; Agarwal, D. S.; Dalela, M.; Mohanty, S.; Sakhuja, R.; Mittal, A.; Chitkara, D. Folate-Targeted Cholesterol-Grafted Lipo-Polymeric Nanoparticles for Chemotherapeutic Agent Delivery. AAPS PharmSciTech 2020, 21 (7), 280. DOI: 10.1208/s12249-020-01812-y.

(8) Apte, A.; Kapoor, M.; Naik, S.; Lubree, H.; Khamkar, P.; Singh, D.; Agarwal, D.; Roy, S.; Kawade, A.; Juvekar, S.; et al. Efficacy of transdermal delivery of liposomal micronutrients through body oil massage on neurodevelopmental and micronutrient deficiency status in infants: results of a randomized placebo-controlled clinical trial. BMC Nutr 2021, 7 (1), 48. DOI: 10.1186/s40795-021-00458-8.

(9) Uddin, M. J.; Werfel, T. A.; Crews, B. C.; Gupta, M. K.; Kavanaugh, T. E.; Kingsley, P. J.; Boyd, K.; Marnett, L. J.; Duvall, C. L. Fluorocoxib A loaded nanoparticles enable targeted visualization of cyclooxygenase-2 in inflammation and cancer. Biomaterials 2016, 92, 71–80. DOI: 10.1016/j.biomaterials.2016.03.028.

(10) Agarwal, S.; Maekawa, T. Nano delivery of natural substances as prospective autophagy modulators in glioblastoma. Nanomedicine 2020, 29, 102270. DOI: 10.1016/j.nano.2020.102270.

(11) Uddin, M. J.; Crews, B. C.; Blobaum, A. L.; Kingsley, P. J.; Gorden, D. L.; McIntyre, J. O.; Matrisian, L. M.; Subbaramaiah, K.; Dannenberg, A. J.; Piston, D. W.; et al. Selective visualization of cyclooxygenase-2 in inflammation and cancer by targeted fluorescent imaging agents. Cancer Res 2010, 70 (9), 3618–3627. DOI: 10.1158/0008-5472.CAN-09-2664.

(12) Uddin, M. J.; Lo, J. H.; Oltman, C. G.; Crews, B. C.; Huda, T.; Liu, J.; Kingsley, P. J.; Lin, S.; Milad, M.; Aleem, A. M.; et al. Discovery of a Redox-Activatable Chemical Probe for Detection of Cyclooxygenase-2 in Cells and Animals. ACS Chem Biol 2022, 17 (7), 1714–1722. DOI: 10.1021/acschembio.1c00961.

(13) Uddin, M. J.; Crews, B. C.; Ghebreselasie, K.; Daniel, C. K.; Kingsley, P. J.; Xu, S.; Marnett, L. J. Targeted imaging of cancer by fluorocoxib C, a near-infrared cyclooxygenase-2 probe. J Biomed Opt 2015, 20 (5), 50502. DOI: 10.1117/1.JBO.20.5.050502.

(14) Uddin, M. J.; Crews, B. C.; Ghebreselasie, K.; Marnett, L. J. Design, synthesis, and structure-activity relationship studies of fluorescent inhibitors of cycloxygenase-2 as targeted optical imaging agents. Bioconjug Chem 2013, 24 (4), 712–723. DOI: 10.1021/bc300693w.

(15) Uddin, M. J.; Crews, B. C.; Huda, I.; Ghebreselasie, K.; Daniel, C. K.; Marnett, L. J. Trifluoromethyl fluorocoxib a detects cyclooxygenase-2 expression in inflammatory tissues and human tumor xenografts. ACS Med Chem Lett 2014, 5 (4), 446–450. DOI: 10.1021/ml400485g.

(16) Ra, H.; Gonzalez-Gonzalez, E.; Uddin, M. J.; King, B. L.; Lee, A.; Ali-Khan, I.; Marnett, L. J.; Tang, J. Y.; Contag, C. H. Detection of non-melanoma skin cancer by in vivo fluorescence imaging with fluorocoxib A. Neoplasia 2015, 17 (2), 201–207. DOI: 10.1016/j.neo.2014.12.009.

(17) Cekanova, M.; Uddin, M. J.; Bartges, J. W.; Callens, A.; Legendre, A. M.; Rathore, K.; Wright, L.; Carter, A.; Marnett, L. J. Molecular imaging of cyclooxygenase-2 in canine transitional cell carcinomas in vitro and in vivo. Cancer Prev Res (Phila) 2013, 6 (5), 466–476. DOI: 10.1158/1940-6207.CAPR-12-0358.

(18) Cekanova, M.; Uddin, M. J.; Legendre, A. M.; Galyon, G.; Bartges, J. W.; Callens, A.; Martin-Jimenez, T.; Marnett, L. J. Single-dose safety and pharmacokinetic evaluation of fluorocoxib A: pilot study of novel cyclooxygenase-2-targeted optical imaging agent in a canine model. J Biomed Opt 2012, 17 (11), 116002. DOI: 10.1117/1.JBO.17.11.116002.

(19) Udddin, M. J.; Moore, C. E.; Crews, B. C.; Daniel, C. K.; Ghebreselasie, K.; McIntyre, J. O.; Marnett, L. J.; Jayagopal, A. Fluorocoxib A enables targeted detection of cyclooxygenase-2 in laser-induced choroidal neovascularization. J Biomed Opt 2016, 21 (9), 90503. DOI: 10.1117/1.JBO.21.9.090503.

(20) Doonze, O.; Picard, D. RNA interference in mammalian cells using siRNAs synthesized with T7 RNA polymerase. Nucleic Acids Res 2002, 30 (10), e46. DOI: 10.1093/nar/30.10.e46 From NLM Medline.

(21) Udddin, M. J.; Crews, B. C.; Xu, S.; Ghebreselasie, K.; Daniel, C. K.; Kingsley, P. J.; Banerjee, S.; Marnett, L. J. Antitumor Activity of Cytotoxic Cyclooxygenase-2 Inhibitors. ACS Chem Biol 2016, 11 (11), 3052–3060. DOI: 10.1021/acschembio.6b00560.

(22) Udddin, M. J.; Marnett, L. J. Synthesis of 5-and 6-carboxy-X-rhodamines. Org Lett 2008, 10 (21), 4799–4801. DOI: 10.1021/ol801904k.

(23) Udddin, M. J.; Smithson, D. C.; Brown, K. M.; Crews, B. C.; Connelly, M.; Zhu, F.; Marnett, L. J.; Guy, R. K. Podophyllotoxin analogues active versus Trypanosoma brucei. Bioorg Med Chem Lett 2010, 20 (5), 1787–1791. DOI: 10.1016/j.bmcl.2010.01.009.

(24) Sahah, H.; Pang, L.; Wang, H.; Shu, D.; Qian, S..; Sathish, V. Growth inhibitory and anti-metastatic activity of epithelial cell adhesion molecule targeted three-way junctional delta-5-desaturase siRNA nanoparticle for breast cancer therapy. Nanomedicine 2020, 30, 102298. DOI: 10.1016/j.nano.2020.102298.

(25) Neil, J. R.; Johnson, K. M.; Nemenoff, R. A.; Schiemann, W. P. Cox-2 inactivates Smad signaling and enhances EMT stimulated by TGF-beta through a PGE2-dependent mechanisms. Carcinogenesis 2008, 29 (11), 2227–2235. DOI: 10.1093/carcin/bgn202 From NLM Medline.

