## Supporting Information BioRxiv1 for "Polymeric Micellar Nanoparticles Enable Image-guided Drug Delivery in Solid Tumors"

**Solvents and Reagents.** We used HPLC grade solvents that were purchased from Fisher (Pittsburgh, PA), and ACS grade starting materials and NMR solvents obtained from the Millipore-Sigma (St. Louis, MO).

**Chromatography.** Standard grade Sorbent silica gel was used for flash column chromatography and a reversed-phase HPLC (Waters 1525 Binary HPLC with 2489 UV/Vis Detector) using a C18 column (Phenomenex Luna 00G-4252-P0- AX) with gradient between water and acetonitrile (20% to 80% acetonitrile over 10 minutes followed by 60% to 100% acetonitrile over 25 minutes, then held for 5 additional minutes) was used for purification of products.

**Spectroscopic Characterization of Small Molecules.** A Bruker 600 MHz instrument was used for $^1$H NMR and $^1$H COSY experiments of all small molecules dissolved in DMSO-$d_6$. The acquired data was processed using a squared sine-bell window function and symmetrized the data for display. The mass spectroscopic analysis was performed on an LTQ Orbitrap high resolution mass spectrometer (Thermo) interfaced to a Waters Acquity UPLC system (Waters Corp., Milford, MA). The mass spectra were acquired in positive or negative ion mode over a precursor ion scan range of m/z 150 to 2000 and the data acquisition was done using Thermo-Finnigan Xcalibur version 2.0.7 and LTQ Orbitrap MS version 2.5.5.

**Spectroscopic Characterization of Polymers.** A Brüker 400 MHz spectrometer was used for $^1$H NMR spectroscopic analysis of all polymers dissolved in $CDCl_3$. The number average molecular weights (Mn), and polydispersities (PDI) were measured using gel permeation chromatography (GPC) with an Agilent Technologies system (Santa Clara, CA, USA). For the GPC analysis, dimethylformamide (DMF) with 0.1 M LiBr was used as the mobile phase at 60 °C, passing through three serial Tosoh Biosciences TSKGel Alpha columns (Tokyo, Japan). To calculate the absolute molecular weights using GPC, serial dilutions (ranging from 10 mg/mL to 0.25 mg/mL) were analyzed on a digital refractometer to determine the polymers' refractive index increment (dn/dc).

**Preclinical Optical Imaging System.** We used a Xenogen IVIS 200 Optical Imaging System for fluorescence imaging of NU/J mice harboring orthotopic breast tumors. The imaging system was configured with a DsRed filter spanning 570–615 nm with a 20 cm field of view at 20 microns resolution.[1] We anesthetized tumor bearing mice with 2% isoflurane (Piramal Critical Care, Bethlehem PA) and subcutaneously (s.c.) injected them with CA-FQ-NPs solution at a dose of 1 mg/kg of each compound. The instrument was designed for simultaneous fluorescence and bright-field imaging.[2]

**Optical Imaging of Orthotopic Breast Tumors.** We dosed female NU/J mice (n = 6 animals/group) bearing orthotopic breast tumors (800-900 $mm^3$) with CA-FQ-NPs (1 mg/kg CA or FQ, s.c.) dissolved in DPBS. At 49 h post-injection, we lightly anesthetized dosed animals with 2% isoflurane and imaged them using a Xenogen IVIS 200 Optical Imaging Instrument with a DsRed filter at a depth of 1.5 cm and an exposure of 1 sec.[2]

**Comparison of Fluorescence Intensities**. Using an ImageJ software, we analyzed the static images obtained from the optical imaging of tumor bearing animals (n = 6 animals/group), where regions of interest (ROIs) were created. We measured the fluorescence intensities within the ROIs and compared the intensities in tumorous versus non-tumorous breasts.

**Measurement of Fluorescence Intensity.** *In vivo* distribution of CA-FQ-NPs was evaluated in female NU/J mice (n = 6) bearing orthotopic breast tumors. At 49 h post-injection of CA-FQ-NPs (100 μl, 290 μg). After imaging, we sacrificed mice overdose. Following dissection, tumorous breast, brain, liver, lung, kidney, non-tumorous breast was collected. The organs were imaged by a Xenogen IVIS 200 Optical Imaging Instrument. Fluorescence intensity (photons/sec) in static image of each organ was measured using an ImageJ software.[3]

**Measurement of Signal-to-Noise Ratio**. The signal-to-noise ratios was measured by dividing the mean fluorescence intensity of tumorous breast with the mean fluorescence intensity of control breast within the respective ROIs measured in photons/sec using an ImageJ software.[4]

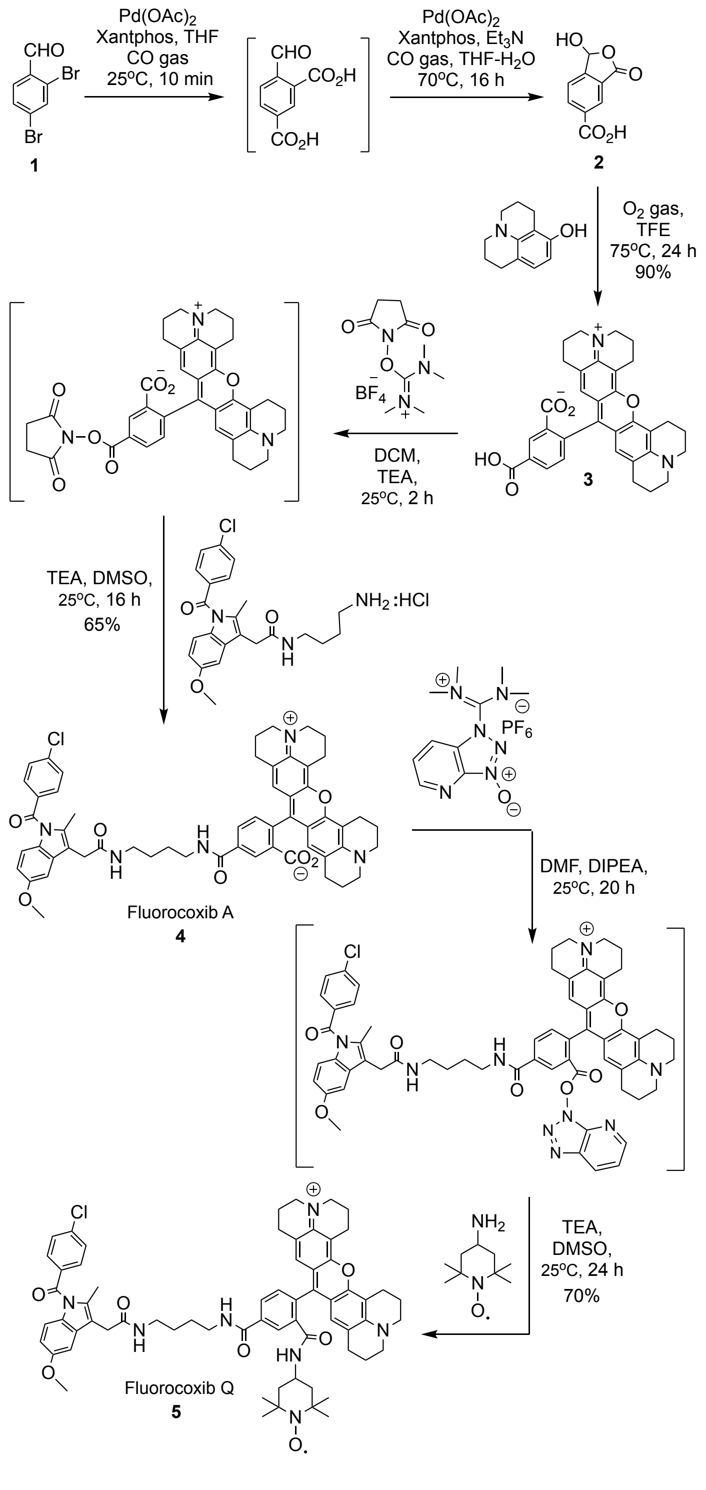


**Scheme S1.** Chemical synthesis of fluorocoxib Q.

**6-Carboxy-3-hydroxyisobenzofuranone (2).** Degassed tetrahydrofuran (25 mL) was added to a mixture of 2,4-dibromobenzaldehyde (**1**, 710 mg, 2.69 mmol), $Pd(OAc)_2$ (30 mg, 135 μmol) and

Xantphos (4,5-Bis(diphenylphosphino)-9,9-dimethylxanthene, 156 mg, 269 μmol) under argon in a 100 mL round bottomed flask. The reaction flask was evacuated *in vacuo* and back filled with gaseous carbon monoxide (CO). The CO-gas was bubbled through the solution for 10 min at 25°C, followed by addition of water (2.5 mL) and triethylamine (TEA, 1.9 mL). The reaction mixture was stirred 16 h at 70°C under CO atmosphere. The reaction mixture was allowed to cooldown to room temperature and the solvent was evaporated *in vacuo* to give a residue. To the residue was added DCM (50 mL) and water (75 mL). The pH was adjusted to 10 using aqueous 2N KOH, and the product was extracted with water (3 x 50 mL). The aqueous layers were acidified with 1N HCl at pH 2 with and the product extracted with EtOAc (3 x 150 mL). The organic layers were combined, dried over anhydrous $Na_2SO_4$, filtered, concentrated and performed a silica gel column chromatography using $CHCl_3$:MeOH:$NH_4OH$ (35:7:1) to afford 6-carboxy-3-hydroxyisobenzofuranone (**2**) as an off white solid (366 mg, 72%). $^1$H NMR (600 MHz, DMSO-$d_6$) δ 6.75 (s, 1H), 7.81 (d, J = 2.5 Hz, 1H) 8.24 – 8.33 (m, 3H), 13.59 (br s, 1H); HRMS (ESI$^-$) *m/z* for $C_9H_5O_5^-$ [M-H]$^-$ calcd 193.0215; found 193.0211.

**5-Carboxy-X-rhodamine (3).** To a solution of 6-carboxy-3-hydroxyisobenzofuranone (**2**, 116 mg, 0.6 mmol) and 8-hydroxyjulolidine (238 mg, 1.25 mmol) in trifluoroethanol (TFE, 25 mL) was bubbled $O_2$ gas for 10 min at 25°C. The reaction mixture was stirred 24 h at 75°C under $O_2$ atmosphere. The reaction mixture was allowed to cooldown to room temperature and the solvent was evaporated *in vacuo* to afford the crude product, which was purified by a silica gel column chromatography using MeOH:DCM (0% → 60%) to afford 5-carboxy-X-rhodamine (**3**) as a dark red solid (288 mg, 90%). $^1$H NMR (600 MHz, DMSO-$d_6$) δ 1.75-1.78 (m, 4H), 1.96-1.98 (m, 4H), 2.46-2.48 (m, 2H), 2.53-2.55 (m, 2H), 2.89-2.91 (m, 4H), 3.21-3.26 (m, 4H), 3.47-3.52

(m, 4H), 6.19 (s, 2H) 7.21 (d, J = 2.5 Hz, 1H), 8.18 (dd, J = 8.7, 2.5 Hz, 1H) 8.24 – 8.33 (m, 1H). HRMS (ESI$^+$) *m/z* for $C_{33}H_{31}N_2O_5^+$ $(M+H)^+$ calcd 535.2155; found 535.2156.

**Fluorocoxib A (4).** To a stirred solution of 5-ROX-acid (53 mg, 0.1 mmol) in anhydrous dichloromethane (DCM, 5 mL) was added *N,N,N',N'*-tetramethyl-*O*-(*N*-succinimidyl)uronium tetrafluoroborate (TSTU, 30 mg, 0.1 mmol) and triethylamine (TEA, 20 µL). The resultant solution was stirred 2 h at 25°C. The solvent was removed *in vacuo* to give 5-ROX-*N*-succinimidyl ester, which was added to a solution of *N*-(4-aminobutyl)-2-[1-(4-chlorobenzoyl)-5-methoxy-2-methyl-1*H*-indol-3-yl]acetamide hydrochloride (45 mg, 0.1 mmol) in DMSO (10 mL) containing TEA (20 µL). The reaction mixture was stirred 16 h at 25°C. The solvent was removed, and the product was purified by a silica gel column chromatography to give fluorocoxib A (**4**) as a deep blue solid (54 mg, 65%). $^1$H NMR (500 MHz, DMSO-$d_6$) δ 1.42-1.61 (m, 4H), 1.70-1.82 (m, 4H), 1.90-2.01(m, 4H), 2.18 (s, 3H), 2.44-2.46 (m, 4H), 2.85-2.88 (m, 4H), 3.10-3.13 (m, 2H), 3.18-3.29 (m, 10H), 3.46 (s, 2H), 3.72 (s, 3H), 6.09 (s, 2H), 6.68 (dd, *J* = 9, 2 Hz, 1H), 6.95 (d, *J* = 9 Hz, 1H), 7.13 (d, *J* = 1.9 Hz, 1H), 7.25 (d, *J* = 9 Hz, 1H), 7.64 (d, *J* = 8.8 Hz, 2H), 7.67 (d, *J* = 8.8 Hz, 2H), 8.06 (s, 1H), 8.14 (dd, *J* = 9, 2 Hz, 1H), 8.43 (d, *J* = 2 Hz, 1H), 8.70 (br s, 1H). HRMS (ESI$^+$) *m/z* for $C_{56}H_{55}ClN_5O_7^+$ $(M+H)^+$ calcd 944.3712; found 944.3722.

**Fluorocoxib Q (5).** To a stirred solution of fluorocoxib A (**4**, 94 mg, 0.1 mml) in *N,N*-dimethylformamide (DMF, 10 mL) containing *N,N*-diisopropylethylamine (DIPEA, 20 µL) was added *N*-[(dimethylamino)-1*H*-1,2,3-triazolo-[4,5-*b*]pyridin-1-ylmethylene]-*N*-methylmethanaminium hexafluorophosphate *N*-oxide (HATU, 38 mg, 0.1 mmol). The reaction mixture was stirred 20 h at 25°C. The solvent was evaporated *in vacuo* to give a intermediate, which was dissolved in DMSO (5 mL) and added to a solution of TEMPO-amine (17 mg, 0.1 mmol) in DMSO (5 mL) containing TEA (20 µL). The resultant mixture was stirred 24 h at 25°C. The solvent was evaporated, and the product was purified by a silica gel column chromatography ($CHCl_3$/MeOH/$NH_4OH$, 35:7:1, v/v/v) to give the pure fluorocoxib Q as a pale brown solid (76 mg, 70%). $^1$H NMR (600 MHz, DMSO-$d_6$) δ 0.94-0.98 (m, 4H), 1.33-1.38 (m, 16H), 1.60-1.64 (m, 4H), 1.86-1.89 (m, 4H), 2.15-2.18 (m, 2H), 2.34 (s, 3H), 2.57-2.59 (m, 3H), 2.91-2.93 (m, 2H), 3.10-3.18 (m, 8H), 3.21-3.26 (m, 6H), 3.60 (s, 2H), 3.84 (s, 3H), 5.78 (s, 2H), 6.75-6.77 (m, 1H), 7.01-7.02 (m, 1H), 7.22 (s, 1H), 7.29-7.31 (m, 1H), 7.74-7.77 (m, 4H), 7.78-7.81 (m, 1H), 8.15-8.19 (m, 2H), 8.26-8.27 (m, 1H), 8.81-8.84 (br s, 1H). HRMS (ESI$^+$) calcd for $C_{65}H_{72}ClN_7O_7^{\bullet+}$ ($M^{\bullet+}$) 1097.5176; found 1097.6364.

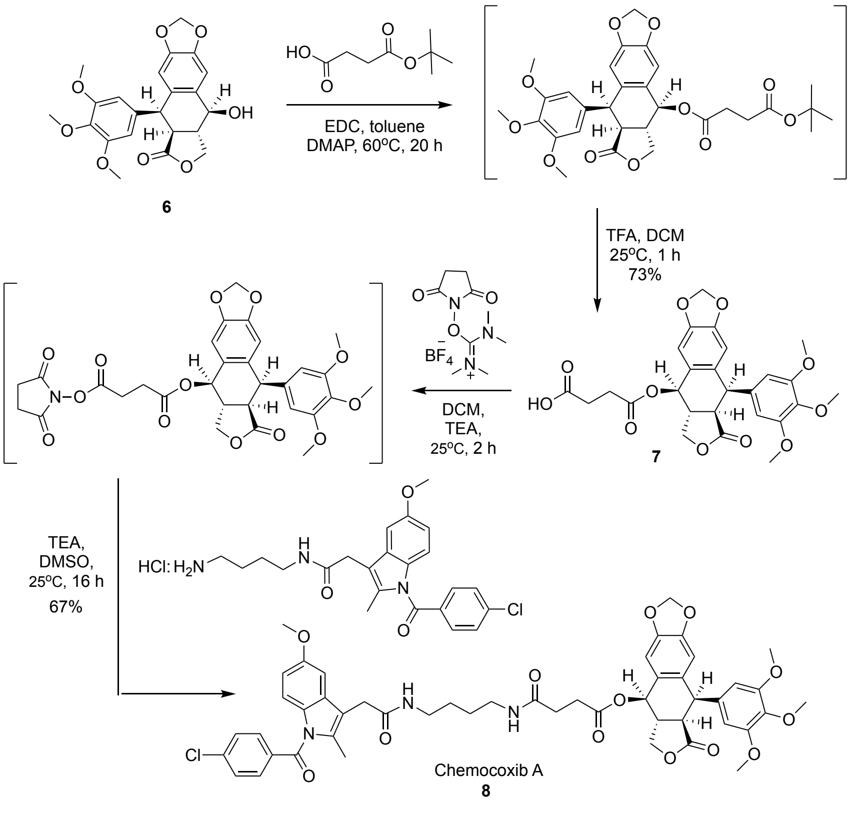


**Scheme S2.** Chemical synthesis of Chemocoxib A.

**Succinylpodophyllotoxin (7).** To a stirred solution of podophyllotoxin (4.14 g, 10 mmol) in toluene (250 mL) was added 4-(*tert*-butoxy)-4-oxobutanoic acid (5.22 g, 30 mmol), 4-dimethylaminopyridine (DMAP, 1 mg) at 25 ºC. The resultant mixture was stirred for 20 h at 60 ºC. The reaction mixture was allowed to cooldown to room temperature. The solvent was removed *in vacuo* and water (100 mL) was added. The product was extracted with EtOAc (3 X 75 mL). The combined organic layer was dried over anhydrous $Na_2SO_4$ and the solvent was evaporated *in vacuo* to give a residue, which was dissolved in DCM (5 mL). To a stirred solution of the residue was added trifluoroacetic acid (TFA, 10 mL) and the reaction mixture was stirred for 1 h at 25 °C. The solvent was removed *in vacuo* to afford the crude product, which was purified using a silica gel column chromatography ($CHCl_3$/MeOH/$NH_4OH$, 35:7:1 v/v/v) to give succinylpodophyllotoxin (**7**) as an off white solid (180 mg, 73%). $^1$H NMR (500 MHz, DMSO-$d_6$) δ 2.10 (d, *J* = 5.4 Hz, 1H), 2.32 (dd, *J* = 12.6, 4.8 Hz, 1H), 2.35-2.37 (m, 1H), 3.52 (s, 3H, OMe), 3.65 (dd, *J* = 10, 7.5 Hz, 2H), 3.82 (s, 6H), 4.15 (t, *J* = 7.9 Hz, 2H), 4.26 (t, *J* = 7.9 Hz, 2H), 4.51 (d, *J* = 5.4 Hz, 1H), 5.89 (s, 2H), 6.00 (s, 2H), 6.67 (s, 1H), 6.94 (s, 1H), 12.15 (s, 1H). HRMS ($ESI^-$) *m/z* for $C_{26}H_{25}O_{11}$ $(M-H)^-$ calcd 513.1475; found 513.1478.

**Chemocoxib A (8).** To a stirred solution of succinylpodophyllotoxin (**10**, 512 mg, 1 mmol) in dichloromethane (10 mL) was added *N,N,N',N'*-tetramethyl-O-(*N*-succinimidyl)uronium tetrafluoroborate (301 mg, 1 mmol) and triethylamine (TEA, 10 mg) at 25 ºC. The reaction mixture was stirred 2 h at 25 ºC. Then the solvent was removed *in vacuo* to give a residue, which was added to a stirred solution of *N*-(4-aminobutyl)-2-(1-(4-chlorobenzoyl)-5-methoxy-2-methyl-1*H*-indol-3-yl)acetamide hydrochloride and triethylamine in DMSO (10 mL). The reaction mixture was stirred 16 h at 25 ºC. The solvent was removed *in vacuo* and the crude product was purified using a silica gel column chromatography ($CHCl_3$/MeOH/$NH_4OH$, 35 : 7 : 1, v/v/v) to give chemocoxib A (**8**) as a pale yellow solid (618 mg, 67 %).$^1$H NMR (500 MHz, DMSO-$d_6$) δ 1.34-1.36 (m, 4H), 3.24 (s, 3H), 3.32-2.36 (m, 4H), 3.65-3.67 (m, 4H), 2.65-2.67 (m, 1H), 3.35-3.37 (m, 1H), 3.45 (s, 2H), 3.64 (s, 3H), 3.66 (s, 6H), 3.75 (s, 3H), 4.14-4.18 (m, 1H), 4.34-4.38 (m, 2H), 4.54-4.57 (m, 1H), 6.00-6.02 (m, 2H), 6.34 (s, 2H), 6.60 (s, 1H), 6.68 (dd, $J$ = 9, 2.5 Hz, 1H), 6.90 (d, $J$ = 8.5 Hz, 1H), 6.95 (s, 1H), 7.16 (d, $J$ = 2.5 Hz, 1H), 7.64 (d, $J$ = 9 Hz, 2H), 7.66 (d, $J$ = 9 Hz, 2H), 7.98 br (s, 1H) 8.05 (br s, 1H). HRMS (ESI$^+$) *m/z* for $C_{49}H_{51}ClN_3NaO_{13}$ $(M+Na)^+$ calcd 946.3032; found 946.3334.

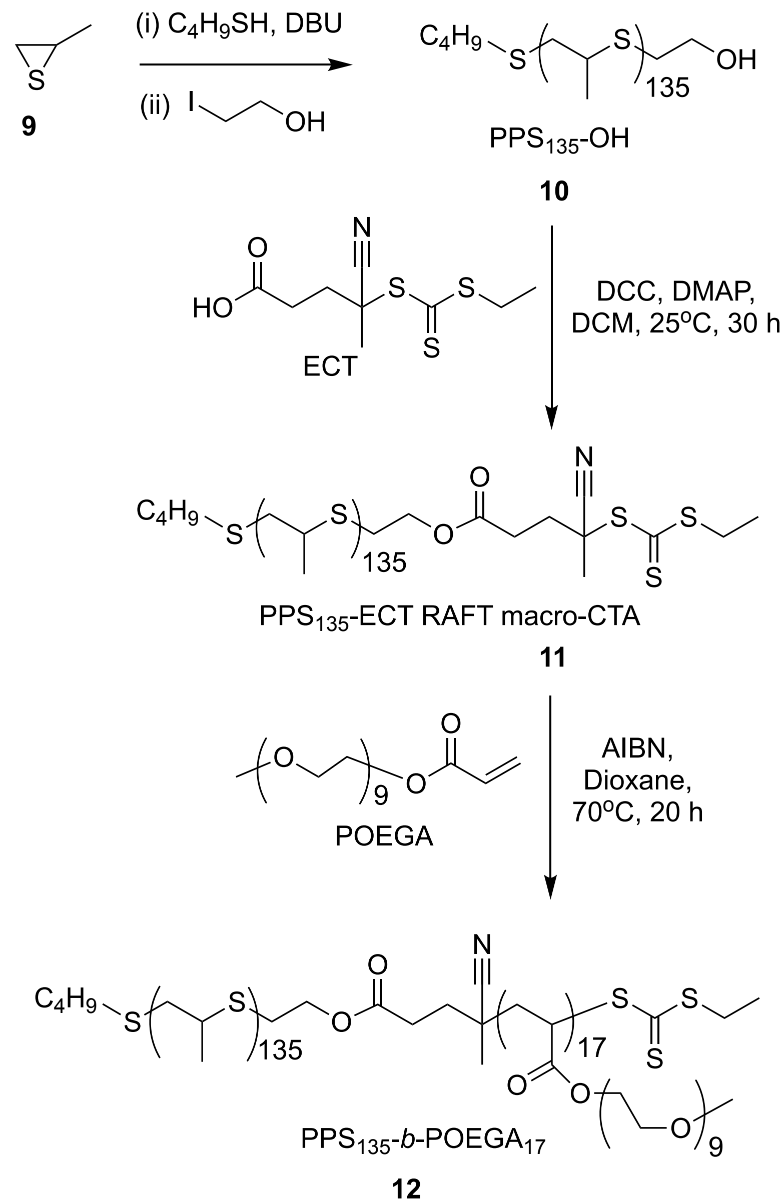


**Scheme S3.** Chemical synthesis of $PPS_{135}$-*b*-$POEGA_{17}$ co-polymer.

**Hydroxyl end functional poly(propylene sulfide) ($PPS_{135}$-OH, 10).** In a nitrogen-flushed 50 mL round-bottom flask, 15 mL of degassed THF and 1-butane thiol (1.0 mmol, 0.070 mL) were added[5, 6]. A degassed solution of 1,8-diazabicyclo[5.4.0]undec-7-ene (DBU) (3 mmol, 0.448 mL) in dry THF (15 mL) was then added dropwise to the flask while maintaining a temperature of 0 ºC, and the mixture was stirred at room temperature for 30 minutes. Afterward, a degassed solution of propylene sulfide (135 mmol, 9.99 g, 10.56 mL) was added, and the mixture was stirred at room temperature for 3 hours. The polymerization process was terminated by adding 2-iodoethanol (5 mmol, 0.85 mL, 0.38 g) and stirring the mixture overnight at room temperature. The following day, the polymer solution was filtered to remove the white precipitated salt, and the crude-filtered polymer solution was precipitated three times in methanol. The viscous polymer was then dried under high vacuum to yield a colorless, viscous polymer. $^1$H NMR (400 MHz; $CDCl_3$, δ): 1.35 (s, C**H$_3$**, **PPS backbone**), 2.52-2.78 (s, -CH, PPS main chain), 2.8-3.1 (s, C**H$_2$, PPS backbone)**, 3.72 (t, C**H$_2$**-OH, PPS terminal end).

**RAFT end functional PPS macro-CTA (PPS$_{135}$-ECT RAFT macro CTA, 11).** A solution of N, N'-Dicyclohexylcarbodiimide (DCC) (0.393 g, 1.5 mmol) was dropwise added to mixture of PPS$_{135}$-OH (3 g, 0.3 mmol), ECT (0.393 g, 1.5 mmol), and 4-Dimethylaminopyridine (DMAP) (0.018 g, 0.15 mmol) in anhydrous DCM (20 mL).[7, 8] The mixture was stirred in the dark for 24 hours at room temperature. After this period, the solution was filtered to remove the precipitated dicyclohexyl urea and then concentrated using rotavap. The resulting viscous polymer was dissolved in DCM and purified by five precipitations in cold methanol. $^1$H NMR (400 MHz; $CDCl_3$, δ): 1.35 (t, 3H, −S−$CH_2$−C$H_3$, ECT), 1.3-1.4 (s, 3H, C**H$_3$, PPS block**), 1.85 (s−C(CN)−C$H_3$, ECT), 2.41−2.67 (m, −C$H_2$−C$H_2$−S, ECT), 2.52-2.78 (broad s, S-CH, PPS backbone), 2.8-3.1 (broad s, 2H, C**H$_2$, PPS backbone**), 3.42 (q, −S−C$H_2$−$CH_3$, ECT), 4.2 (t, -OC**H$_2$**-$CH_2$, PPS end). (PPS$_{135}$-ECT, $M_{n,\,GPC}$= 10,200 g/mol, PDI = 1.13).

**Diblock copolymer (PPS$_{135}$-b-POEGA$_{17}$, 12).** The amphiphilic diblock copolymer PPS$_{135}$-*b*-POEGA$_{17}$ was synthesized through RAFT polymerization of OEGA, using AIBN as initiator and PPS$_{135}$-ECT as the chain transfer agent.[6, 9] In a 25 mL flask, PPS$_{135}$-ECT (1.0 g, 0.1 mmol),

POEGA (1.44 g, 3 mmol), AIBN (1.68 mg, 0.01 mmol), and 5 mL of dioxane were combined. The solution was degassed for 30 minutes before being heated to 70 °C for 24 hours. The following day, the polymerization mixture was cooled to room temperature, precipitated twice in cold diethyl ether, and dried under vacuum overnight at 40 °C to obtain a viscous, yellow-colored polymer. $^1$H NMR (400 MHz; $CDCl_3$, δ): 1.3-1.4 (s, $CH_3$ in PPS block), 1.6-1.9 (s, $CH_2$ and CH backbone protons in POEGA block), 2.5–2.8 (broad s, S-CH, PPS block), 2.8–3.1 (broad s, 2H, $CH_2$, PPS block), 3.3 (s, $OCH_3$, pendent terminal methyl in POEGA), 3.68-3.80 (m, -$OCH_2$-$CH_2$, pendent PEG units in POEGA). ($PPS_{135}$-*b*-$POEGA_{17}$-ECT, $M_{n,\ GPC}$= 18,360 g/mol, PDI = 1.33).

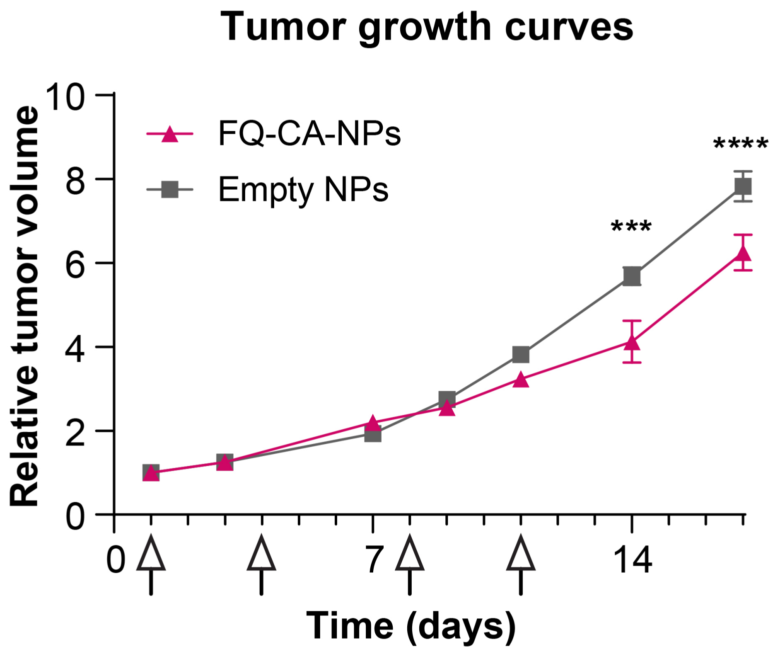


**Figure S1. Therapeutic effect of FQ-CA-NPs in orthotopic triple-negative breast cancer model.** Tumor growth curves for mice with orthotopic MDA-MB-231 tumors treated with FQ-CA-NPs or Empty NPs intravenously (n=4 each, arrows indicate timing of doses). ***: $p<0.001$, ****: $p<0.0001$ by two-way ANOVA comparing the two groups at each timepoint with Šídák's multiple comparisons test. Error bars indicate SEM.

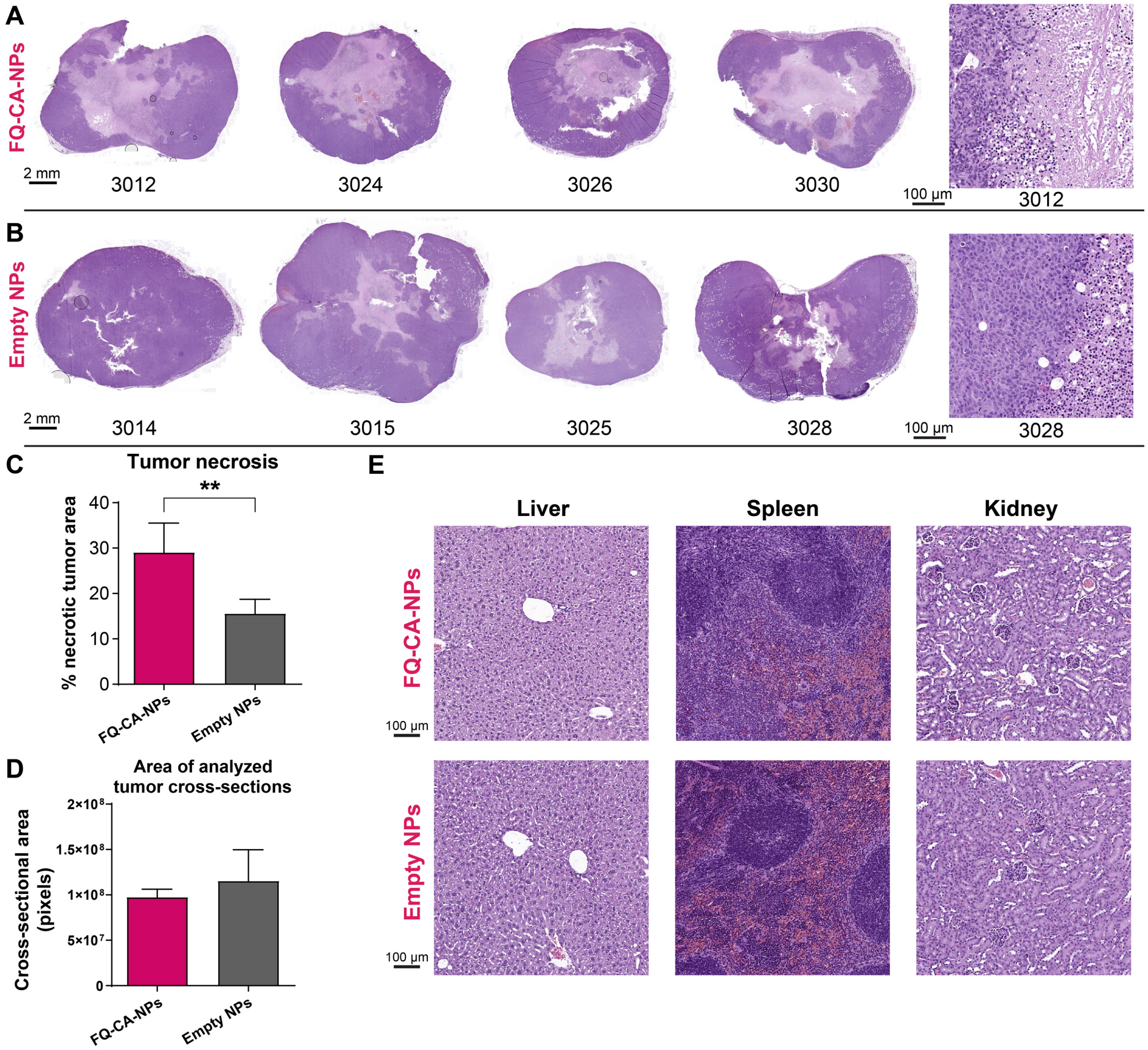


**Figure S2. (A-B)** Representative hematoxylin & eosin (H&E)-stained cross-sections of MDA-MB-231 orthotopic xenograft tumors in mice treated with **(A)** FQ-CA-NPs or **(B)** empty (control) NPs. Numbers below the images indicate mouse ID. Scale bars = 2 mm for full tumor cross-sections shown in panels A and B. Representative higher-magnification H&E images from selected treated tumor specimens are shown at right for each treatment group, with scale bar = 100 µm for these images. **(C)** Comparison of tumor necrotic area in cross-sections of the MDA-MB-231 tumors shown in panels A-B. Two cross-sections were analyzed and averaged per tumor specimen; necrotic area was calculated as a percentage of the total tumor area. **: $p<0.01$ by unpaired, two-tailed t-test. **(D)** Average cross-sectional area of tumor sections used in necrosis analysis. **(E)** Representative images of H&E-stained sections of livers, spleens, and kidneys from mice treated with FQ-CA-NPs vs. Empty NPs; scale bar = 100 µm.

### 8. REFERENCES

(1) Uddin, M. J.; Crews, B. C.; Blobaum, A. L.; Kingsley, P. J.; Gorden, D. L.; McIntyre, J. O.; Matrisian, L. M.; Subbaramaiah, K.; Dannenberg, A. J.; Piston, D. W.; et al. Selective visualization of cyclooxygenase-2 in inflammation and cancer by targeted fluorescent imaging agents. *Cancer Res* **2010**, *70* (9), 3618-3627. DOI: 10.1158/0008-5472.CAN-09-2664 From NLM Medline.

(2) Uddin, M. J.; Crews, B. C.; Ghebreselasie, K.; Marnett, L. J. Design, synthesis, and structure-activity relationship studies of fluorescent inhibitors of cycloxygenase-2 as targeted optical imaging agents. *Bioconjug Chem* **2013**, *24* (4), 712-723. DOI: 10.1021/bc300693w From NLM Medline.

(3) Uddin, M. J.; Crews, B. C.; Huda, I.; Ghebreselasie, K.; Daniel, C. K.; Marnett, L. J. Trifluoromethyl fluorocoxib a detects cyclooxygenase-2 expression in inflammatory tissues and human tumor xenografts. *ACS Med Chem Lett* **2014**, *5* (4), 446-450. DOI: 10.1021/ml400485g From NLM PubMed-not-MEDLINE.

(4) Uddin, M. J.; Lo, J. H.; Oltman, C. G.; Crews, B. C.; Huda, T.; Liu, J.; Kingsley, P. J.; Lin, S.; Milad, M.; Aleem, A. M.; et al. Discovery of a Redox-Activatable Chemical Probe for Detection of Cyclooxygenase-2 in Cells and Animals. *ACS Chem Biol* **2022**, *17* (7), 1714-1722. DOI: 10.1021/acschembio.1c00961 From NLM Medline.

(5) Bezold, M. G.; Hanna, A. R.; Dollinger, B. R.; Patil, P.; Yu, F.; Duvall, C. L.; Gupta, M. K. Hybrid Shear-Thinning Hydrogel Integrating Hyaluronic Acid with ROS-Responsive Nanoparticles. *Advanced Functional Materials* **2023**, *33* (31), 2213368. DOI: https://doi.org/10.1002/adfm.202213368 (acccessed 2024/01/12).

(6) Uddin, M. J.; Werfel, T. A.; Crews, B. C.; Gupta, M. K.; Kavanaugh, T. E.; Kingsley, P. J.; Boyd, K.; Marnett, L. J.; Duvall, C. L. Fluorocoxib A loaded nanoparticles enable targeted visualization of cyclooxygenase-2 in inflammation and cancer. *Biomaterials* **2016**, *92*, 71-80. DOI: https://doi.org/10.1016/j.biomaterials.2016.03.028.

(7) Gupta, M. K.; Meyer, T. A.; Nelson, C. E.; Duvall, C. L. Poly(PS-b-DMA) micelles for reactive oxygen species triggered drug release. *Journal of Controlled Release* **2012**, *162* (3), 591-598. DOI: 10.1016/j.jconrel.2012.07.042.

(8) Gupta, M. K.; Martin, J. R.; Dollinger, B. R.; Hattaway, M. E.; Duvall, C. L. Thermogelling, ABC Triblock Copolymer Platform for Resorbable Hydrogels with Tunable, Degradation-Mediated Drug Release. *Advanced Functional Materials* **2017**,  (27), 1704107.

(9) Gupta, M. K.; Martin, J. R.; Werfel, T. A.; Shen, T.; Page, J. M.; Duvall, C. L. Cell Protective, ABC Triblock Polymer-Based Thermoresponsive Hydrogels with ROS-Triggered Degradation and Drug Release. *Journal of the American Chemical Society* **2014**, *136* (42), 14896-14902. DOI: 10.1021/ja507626y.
